# CREB/CRTC2-Induced GPR65 Orchestrates Th17 Metabolic Fitness and Pathogenic Effector Function

**DOI:** 10.64898/2026.03.27.714848

**Authors:** Si Cave, Dhruv Patel, Qiyuan Yang, Yuqiong Liang, Jack Swanson, Kolby Versage, Ijeoma Chikezie, Jose E Carra Hernandez, Caitlyn Marie Ybanez, Ezra Wiater, Li V. Yang, Ye Zheng, Jeniffer B. Hernandez

## Abstract

GPR65 has been shown to be a critical regulator of Th17 cell pathogenicity. Loss of GPR65 in mice results in a decrease in Th17 cells and reduced susceptibility to a mouse model of multiple sclerosis. The CREB/CRTC2 pathway has emerged as an important regulator of immune function. We have previously shown that the CREB/CRTC2 pathway modulates autoimmune disease by promoting differentiation of Th17 cells. In this study we performed RNA-seq to identify Th17 genes regulated by the CREB/CRTC2 pathway. Our RNA-seq analysis led us to uncover the first mechanism of regulation of the orphan receptor GPR65 by the CREB/CRTC2 pathway. We show that GPR65 is a target of the CREB/CRTC2 pathway through expression studies and chromatin immunoprecipitation. In addition, we show that targeting GPR65 with small molecules alters the expression of IL-17A. Understanding the regulation of GPR65 will be crucial in developing small molecules to treat patients with Th17 cell-mediated disorders.

## INTRODUCTION

Autoimmune diseases represent a major global health burden, affecting millions of individuals worldwide. More than 100 autoimmune conditions have been identified, including inflammatory bowel disease (IBD), which comprises Crohn’s disease (CD) and ulcerative colitis (UC), systemic lupus erythematosus, rheumatoid arthritis, psoriasis, and multiple sclerosis. These disorders arise when the immune system mistakenly targets healthy tissues, leading to chronic inflammation and progressive tissue damage. Approximately 5–8% of the U.S. population is affected by at least one autoimmune condition, with women comprising nearly 75% of diagnosed cases, which is likely due to complex hormonal, genetic, and environmental factors. Although therapeutic strategies such as immunosuppressive agents can alleviate symptoms and slow disease progression, they often carry substantial side effects and may be insufficient for many patients. Improved understanding of the cellular and molecular mechanisms driving autoimmunity is therefore essential for the development of safer and more effective treatments.

Naïve CD4⁺ T cells differentiate into distinct T-helper subsets, including Th1, Th2, Treg, and Th17 cells depending on the cytokine and costimulatory signals received during activation [1]. Each lineage contributes uniquely to host defense and immune regulation. Among these, T helper 17 (Th17) cells have emerged as key drivers of autoimmune pathology. Aberrant Th17 differentiation and expansion are strongly associated with multiple sclerosis, rheumatoid arthritis, psoriasis, and other chronic inflammatory diseases [2,3]. Th17 cells exert their effects largely through the secretion of IL-17A and IL-17F, along with additional inflammatory mediators such as IL-21, IL-22, and granulocyte–macrophage colony-stimulating factor, which collectively promote neutrophil and macrophage recruitment [4–6]. The defining expression of IL-17A and IL-17F enables robust in vitro modeling of Th17 differentiation and provides a tractable system for dissecting signaling pathways relevant to autoimmune disease.

Th17 cells can differentiate into either non-pathogenic or pathogenic subsets depending on the surrounding cytokine milieu. Non-pathogenic Th17 cells arise under TGF-β + IL-6 conditions [7], whereas cytokines such as IL-1β and IL-23 promote the emergence of highly pro-inflammatory pathogenic Th17 cells [8]. Because our study focuses on signaling pathways active within inflammatory Th17 effector states, we polarized cells under pathogenic Th17 conditions, which generate a stable, IL-23-responsive effector population with robust IL-17A production. These conditions provide a physiologically relevant context for dissecting CREB/CRTC2-dependent regulatory mechanisms and downstream functional consequences. Throughout this study, in vitro assays are used to assess Th17 differentiation, whereas in vivo EAE experiments evaluate Th17 pathogenicity, which reflects their disease-driving function.

The cyclic AMP response element–binding protein (CREB) is a transcription factor that binds cAMP response elements to regulate gene expression [7]. CREB activity contributes to T-cell maturation, survival, and proliferation, particularly under pro-inflammatory conditions [6,9–11]. The CREB-regulated transcription coactivator CRTC2 enhances CREB-dependent transcription when dephosphorylated and translocated to the nucleus [12]. The CREB/CRTC2 pathway has been shown to promote IL-17A expression and augment Th17 differentiation; accordingly, T-cell–specific deletion of CRTC2 confers resistance to experimental autoimmune encephalomyelitis (EAE), a mouse model of multiple sclerosis [13].

G protein-coupled receptors (GPCRs) are involved in many diseases and are currently the direct target of about 40% of modern medical drugs [14]. When ligands bind, a conformational change takes place that allows the GPCR to activate associated guanine nucleotide binding proteins (G-proteins) that begin the cAMP signal transduction pathway [15]. G-protein–coupled receptor 65 (GPR65), also known as TDAG8, is an acid-sensing member of the G2A receptor family that increases cAMP production under acidic conditions, and in vivo studies show that loss of GPR65 markedly reduces cAMP levels [16]. Using single-cell RNA-seq to examine cellular states by cytokine production, Gaublomme et al. show that whole body GPR65 knock out (KO) mice exhibit a decrease in Th17 cell differentiation and consequently resistance to EAE [2]. This suggests that GPR65 is required for Th17 cell differentiation. Hence, identifying therapeutic targets of GPR65 could aid in ameliorating autoimmune diseases in which Th17 cells are elevated, such as in multiple sclerosis.

Recent work suggests that GPR65 contributes to the regulation of Th17 cell functional programs, including pathways associated with cellular metabolism [2, 17–18]. Th17 cells rely on glycolysis, glutamine metabolism, fatty acid synthesis, and mTOR signaling to sustain their inflammatory phenotype. Our findings demonstrate that inhibiting GPR65, either through genetic knockout or pharmacological antagonism, disrupts Th17 differentiation, reduces IL-17 production, and diminishes metabolic activity. Together, these results highlight GPR65 as a critical regulator of Th17 cell function and identify it as a promising molecular target for modulating pathogenic inflammation.

## METHODS

### Mice

C57BL/6 (B6) mice were purchased from Jackson Laboratory. CRTC2^⁻/⁻^ mice were generated as previously described [19]. All mice were on a C57BL/6 background, 6–12 weeks of age, and sex-matched for each experiment. Mice were housed under specific pathogen-free conditions. All studies were approved by the IACUC at the Salk Institute, the University of California, Irvine and the University of La Verne. GPR65^-/-^ mice were generated as previously described [20].

### Reagents

Primers were obtained from Integrated DNA Technologies. Recombinant cytokines (TGF-β and IL-6) were from PeproTech; IL-23 from R&D Systems. Anti-CD3, anti-CD28, anti-IL-4, and anti-IFN-γ antibodies were from BioXCell. IL-17A was purchased from BioLegend. Forskolin (FSK) and PGE₂ were from Sigma-Aldrich. BTB09089 was obtained from MolPort; Zinc62678696 from Enamine. Seahorse assay kits were purchased from Agilent.

### CD4⁺ T Cell Isolation and Differentiation in vitro

CD4⁺ T cells were isolated as previously described [10]. Briefly, splenocytes were enriched using the Dynabeads FlowComp Mouse CD4 Kit (Life Technologies) or EasySep™ Mouse CD4⁺ T Cell Isolation Kit (STEMCELL Technologies). For Fig. 1d, bulk CD4⁺ and CD8⁺ T cells were isolated using EasySep™ kits (STEMCELL Technologies). Naïve CD4⁺ T cells (CD62L^hi CD44^lo CD25⁻) were purified by FACS (FACSAria or InFlux; BD Biosciences).

**Figure 1.**
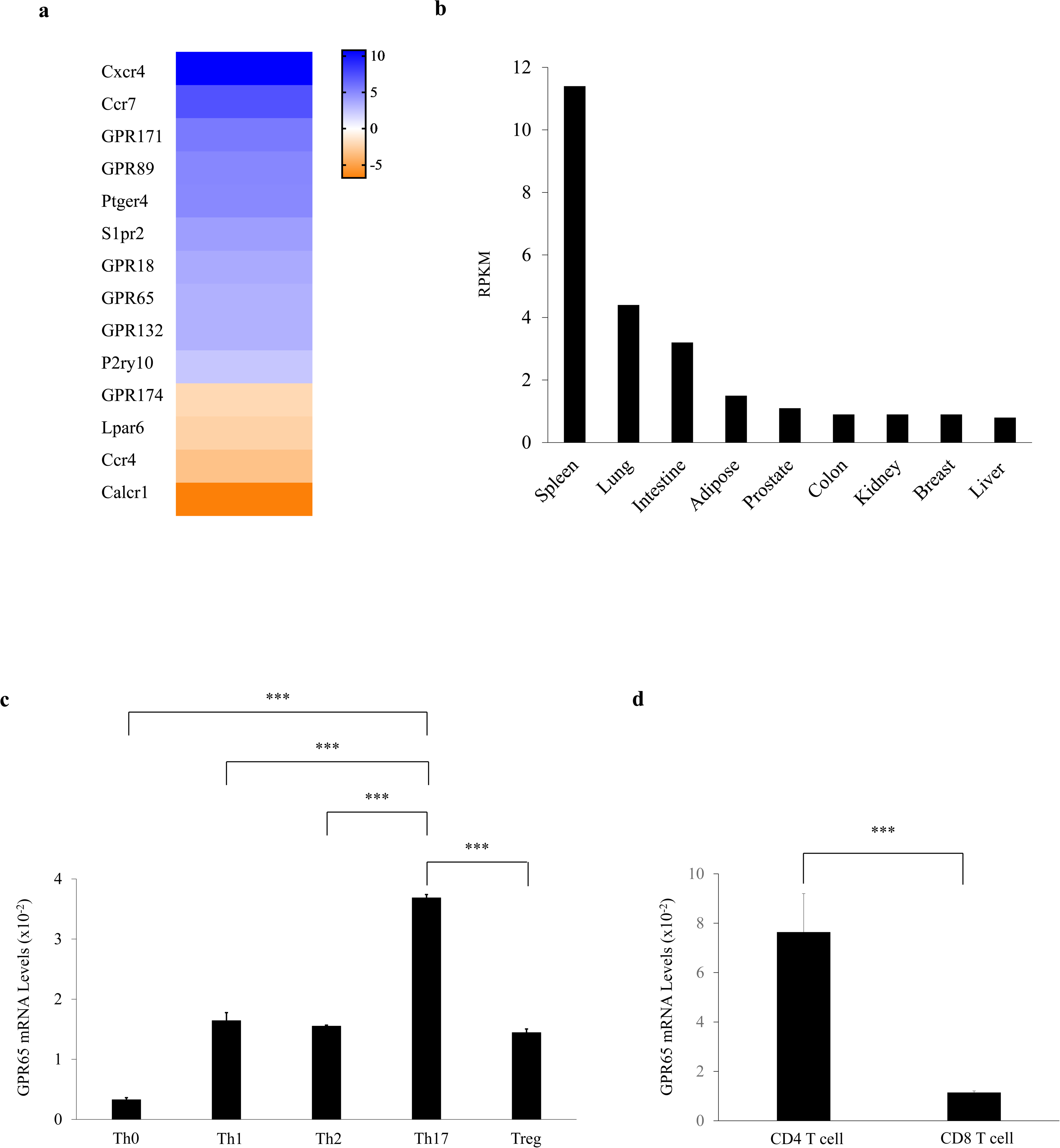
Characterization of GPR65. (a) Heatmap of GPCRs with significant differential expression in Th17 cells treated with forskolin (P < 0.05). Forskolin was added for 1 hour on day 4 of Th17 differentiation prior to RNA extraction. (b) GTEx RNA-seq data showing GPR65 expression (RPKM) across human tissues. (c) GPR65 mRNA levels in naïve CD4⁺ T cells differentiated under Th1, Th2, Treg, or Th17 conditions. Th0 indicates non-polarized cells. Bar graphs represent mean ± s.e.m., n = 3 independent experiments. (d) GPR65 mRNA levels in CD4⁺ versus CD8⁺ T cells. CD4⁺ and CD8⁺ T cells were isolated using the EasySep™ Mouse CD4⁺ or CD8⁺ Isolation Kits (STEMCELL Technologies). Bar graphs represent mean ± s.e.m., n = 3 independent experiments. Statistical analyses were performed using unpaired Student’s t-test. Significance was defined as P < 0.05 (*P < 0.05; **P < 0.005; ***P < 0.0005).

Cells were cultured in Click’s medium (Irvine Scientific) supplemented with 2 mM L-glutamine, 100 U/mL penicillin, 100 μg/mL streptomycin, 50 μM β-mercaptoethanol, and 10% FCS. Plates were coated with anti-hamster IgG (Vector Labs). Naïve CD4⁺ T cells were activated with plate-bound anti-CD3 (1 μg/mL) and soluble anti-CD28 (0.5 μg/mL).

Differentiation conditions were

- Th17: IL-6 (50 ng/mL), TGF-β (1 ng/mL), IL-23 (5 ng/mL), anti-IFN-γ (10 μg/mL), anti-IL-4 (10 μg/mL).
- Th1: IL-12 (10 ng/mL) + anti-IL-4 (10 μg/mL).
- Th2: IL-4 (10 ng/mL) + anti-IFN-γ (10 μg/mL).
- Treg: TGF-β (2 ng/mL) + IL-2 (50 U/mL).

For proliferation assays, Th17 cells were labeled once with CFSE and then cultured for 3 days before analysis. Compounds (FSK, PGE₂, BTB09089, Zinc62678696) were added for 1 hour on day 4 of Th17 differentiation immediately prior to RNA extraction.

### Human Th17 Cell Source

Human peripheral blood samples were purchased from STEMCELL Technologies. PBMCs were isolated using the Ficoll-Paque density-gradient protocol recommended by the manufacturer. Th17 cells were then enriched using the EasySep™ Human Th17 Cell Enrichment Kit II (STEMCELL Technologies; CD4⁺CXCR3⁻CCR6⁺ enrichment using particle release technology; Cat# 18063) according to the manufacturer’s instructions. Immediately after enrichment, cells were resuspended in complete RPMI and treated with vehicle (DMSO) or Zinc62678696 (30 μM) for 1 hour without additional ex vivo cytokine polarization or activation. Cells were then harvested for RNA extraction.

### RNA-Sequencing

RNA-seq was performed using three biological replicates per condition (n = 3), consistent with accepted standards for bulk RNA-seq differential expression studies. Total RNA was isolated from differentiated CD4⁺ T cells and poly-A–purified using poly-dT agarose magnetic beads. Strand-specific libraries were prepared using the NEBNext Ultra Directional RNA Library Prep Kit for Illumina. Libraries were sequenced on an Illumina MiSeq platform using V2 chemistry with 75 bp paired-end reads.

Read depth was selected based on sample complexity and the required sensitivity for differential expression analysis. FASTQ files were aligned to the mouse reference genome using TopHat2, and differential gene expression was quantified using Cuffdiff. Read alignments (accepted_hits.bam files) were converted to bedGraph format for visualization in the UCSC Genome Browser. Gene-level expression analyses were performed using Partek Flow (Partek Inc.). The RNA-seq dataset generated in this study has been deposited in NCBI Gene Expression Omnibus (GEO) under accession number GSE166496 (https://www.ncbi.nlm.nih.gov/geo/query/acc.cgi?acc=GSE166496).

### Real-Time Quantitative PCR

RNA isolation was performed with Trizol reagent using a standard RNA protocol. RNA quantity was assessed with a NanoQuant Infinite M200 instrument (Tecan Group Ltd). cDNA was generated using reverse transcriptase from the Azura Quant cDNA Synthesis Kit (Azura Genomics) on a PTC200 Thermal Cycler (MJ Research) and T100 PCR Gradient Thermal Cycler (Bio-Rad). Using AzuraQuant Green Fast qPCR Mix LoRox Kit (Azura Genomics), cDNA was quantified on a LightCycler 96 System (Roche Life Science). Gene expression was measured relative to the 18S and GAPDH housekeeping genes.

### Chromatin Immunoprecipitation

Chromatin immunoprecipitation using CREB antisera was performed as previously described [13], with minor modifications. Following Th17 differentiation, cells were treated with vehicle (DMSO) or forskolin (10 µM) for 1 hour prior to fixation to assess forskolin-induced CREB recruitment to the *Gpr65* promoter. Cells were fixed in 1% formaldehyde for 10 minutes at room temperature, and crosslinking was quenched by adding 125 mM glycine for 5 minutes while rocking.

Cells were resuspended in cell lysis buffer (25 mM HEPES pH 7.8, 1.5 mM MgCl₂, 10 mM KCl, 0.3% NP-40, 1 mM DTT) and incubated on ice for 10 minutes. Nuclei were pelleted by centrifugation (5,000 rpm, 5 minutes) and lysed in 800 μL of nuclei lysis buffer (50 mM HEPES pH 7.9, 140 mM NaCl, 1 mM EDTA, 1% Triton X-100, 0.1% sodium deoxycholate, 0.2% SDS) for 10 minutes on ice.

Chromatin was sonicated to approximately 200–500 bp fragments (8 cycles of 15 seconds at setting 8). Lysates were cleared by centrifugation, and supernatants were precleared with Protein A for 6 hours. Precleared chromatin was divided equally, and 1% of each sample was saved as input. For immunoprecipitation, 2 μg of anti-CREB (clone 48H2, Cell Signaling Technology, Cat# 9197) was added overnight at 4°C, followed by incubation with Protein A beads for 2 hours. Normal rabbit IgG (Cell Signaling Technology, Cat# 2729) was used as the isotype control and processed in parallel for both vehicle- and forskolin-treated samples.

Beads were washed once with nuclei lysis buffer, once with nuclei lysis buffer containing 500 mM NaCl, twice with wash buffer (20 mM Tris pH 8.0, 1 mM EDTA, 250 mM LiCl, 0.5% NP-40, 0.5% sodium deoxycholate), and twice with TE buffer. DNA-protein complexes were eluted in 1% SDS, 1 mM EDTA, 10 mM Tris pH 8.1, and crosslinks were reversed overnight at 65°C. Samples were then treated with proteinase K for 2 hours at 65°C, supplemented with glycogen, extracted with phenol–chloroform, and ethanol-precipitated. Purified DNA was resuspended in water and analyzed by quantitative PCR using primers flanking the CRE half-site located −5,526 bp upstream of the *Gpr65* transcriptional start site.

### Flow Cytometry

Differentiated CD4⁺ T cells were harvested and analyzed by flow cytometry. Cells were first incubated with Fixable Viability Dye (Thermo Fisher) for live/dead discrimination according to the manufacturer’s protocol. Data acquisition was performed on a BD LSRFortessa and analyzed using FlowJo software (v10). A standardized gating strategy was applied across all experiments:

1. Forward and side scatter to identify lymphocytes,
2. Singlet discrimination (FSC-H versus FSC-A),
3. Live/dead exclusion, and
4. Gating on CD4⁺ T cells

before downstream functional gates were applied.

For Th1, Treg, and Th17 conditions, intracellular cytokine staining was performed following PMA/ionomycin restimulation in the presence of brefeldin A. Cells were fixed and permeabilized using the eBioscience Foxp3/Transcription Factor Staining Buffer Set. Antibodies used included: anti-CD4 (clone RM4), anti-IFN-γ (clone XMG1.2), anti-Foxp3 (clone 150D), and anti-IL-17A (clone TC11-18H10.1) (all from BioLegend or eBioscience).

Fluorescence-minus-one (FMO) controls and isotype controls were included to define cytokine-positive gates. The gating strategy was identical for wild-type and GPR65⁻/⁻ samples and was confirmed across three independent experiments.

### Retroviral Infection

MSCV-IRES-GFP and MSCV-ACREB-IRES-GFP were transfected into 293T cells along with PsiEco packaging vector and retroviral supernatants were collected as previously described [13]. Naïve CD4⁺ T cells were differentiated into Th17 cells as stated above and infected on days 1 and 2. Fresh cytokines were added after each infection. RNA was then isolated as described above on day 4.

### Seahorse assays

To determine the mitoATP, glycoATP, and oxygen consumption rate (OCR), extracellular acidification (ECAR), naïve CD4+ T-cells cultured under Th17 cells for 5 days, were harvested in XF-RPMI medium (pH 7.4, 37°C). Cells were plated in PDL mini plate (XFp) at concentrations of 4×10^5^ cells per well. The mitoATP, glycoATP, oxygen consumption rate (OCR), extracellular acidification (ECAR) were assessed using ATP Rate Assay, Glycolytic rate assay and Cell Mito Stress Test (Agilent) using Seahorse XFp analyzer (Agilent). Assays were performed according to the protocol provided by Agilent.

### Experimental Autoimmune Encephalomyelitis

To induce EAE, mice were immunized with 100 μg MOG₃₅–₅₅ in CFA using the Hooke Kit™ (Cat# EK-2110). Bordetella pertussis toxin (200 ng) was administered intraperitoneally on days 0 and 2. Beginning on **day 15**, mice received **daily** subcutaneous injections of vehicle or Zinc62678696 (75 mg/kg). Mice were monitored for signs of disease on the indicated days. Clinical disease following EAE induction was assessed using a previously described scale (40) as follows: (0.5) altered gait and/or hunched appearance, [1] limp tail, [2] partial hind limb paralysis, [3] complete hind limb paralysis, [4] complete hind limb paralysis and partial fore limb paralysis, and [5] death.

### Statistical analysis

Statistical analyses were performed using GraphPad Prism or Excel. In vitro data are presented as mean ± s.e.m., and comparisons between two groups were assessed by unpaired two-tailed Student’s t-tests.

Because EAE clinical scores are ordinal and non-normally distributed, daily comparisons were assessed using the Mann–Whitney U test. No multiple comparisons correction was applied, consistent with standard EAE analysis. All EAE experiments were performed twice with consistent results. A p value < 0.05 was considered significant.

## RESULTS

### Characterization of GPR65

We have previously shown that Th17 cell differentiation is partially regulated by the CREB/CRTC2 pathway [13]. To identify additional CREB/CRTC2-responsive targets, we performed RNA-seq on differentiated Th17 cells treated with forskolin (FSK), a potent activator of cAMP signaling that induces CREB/CRTC2 activation. Genes were considered CREB-responsive if they met the criteria of log₂ fold change (log₂FC) > 1.0 and adjusted P < 0.05. Several GPCRs satisfied these thresholds and were therefore classified as candidate CREB-regulated genes (**Fig. 1a**).

Among these, *Gpr65* was prioritized for detailed follow-up because (i) it met statistical criteria for CREB responsiveness, (ii) it demonstrated Th17-enriched expression, and (iii) prior work by Gaublomme et al. identified GPR65 as a critical regulator of Th17 cell pathogenicity in vivo [2]. In addition, tissue-level RNA expression data from the Human Protein Atlas showed markedly elevated GPR65 expression in the human spleen compared with other organs (**Fig. 1b**), further supporting an immune-specific role for this receptor [21].

To determine whether *Gpr65* expression is specialized to Th17 cells, we compared transcript levels across multiple CD4⁺ T-cell subsets (**Fig. 1c**). *Gpr65* was significantly upregulated in Th17 cells compared with Th1, Th2, and Treg subsets. Other CREB-responsive GPCRs that met the same statistical thresholds, including *Gpr18* and *Gpr132*, also exhibited Th17-skewed and CRTC2-dependent expression, but lacked prior evidence linking them to Th17 pathogenicity or autoimmune inflammation; therefore, *Gpr65* was selected for mechanistic follow-up (**Fig. S1**). Finally, *Gpr65* expression was substantially lower in CD8⁺ T cells than in CD4⁺ T cells (**Fig. 1d**), suggesting a CD4⁺ T cell–preferential role.

### CREB binds upstream of the Gpr65 Transcriptional Start Site

CREB dimers bind to specific DNA sequences called cAMP response elements (CRE) sites through dimerization of a leucine zipper and phosphorylation at serine-133, either increasing or decreasing gene expression [19]. The CRE-binding sites are located in enhancer and promoter regions and affect specific genes downstream or upstream, depending on the specific location of the CRE site. CRE sites contain highly conserved nucleotide sequences that are either palindromic (5’-TGACGTCA-3’) or half-site (5’-TGACG-3’ or 5’-CGTCA-3) [19, 22]. The *Gpr65* gene contains 4 half site CREs at -5,526, -8,380, -8,597, and -10,674 relative to the transcriptional start site (TSS) suggesting that CREB may indeed transcriptionally regulate *Gpr65* (**Fig. 2a**).

**Figure 2.**
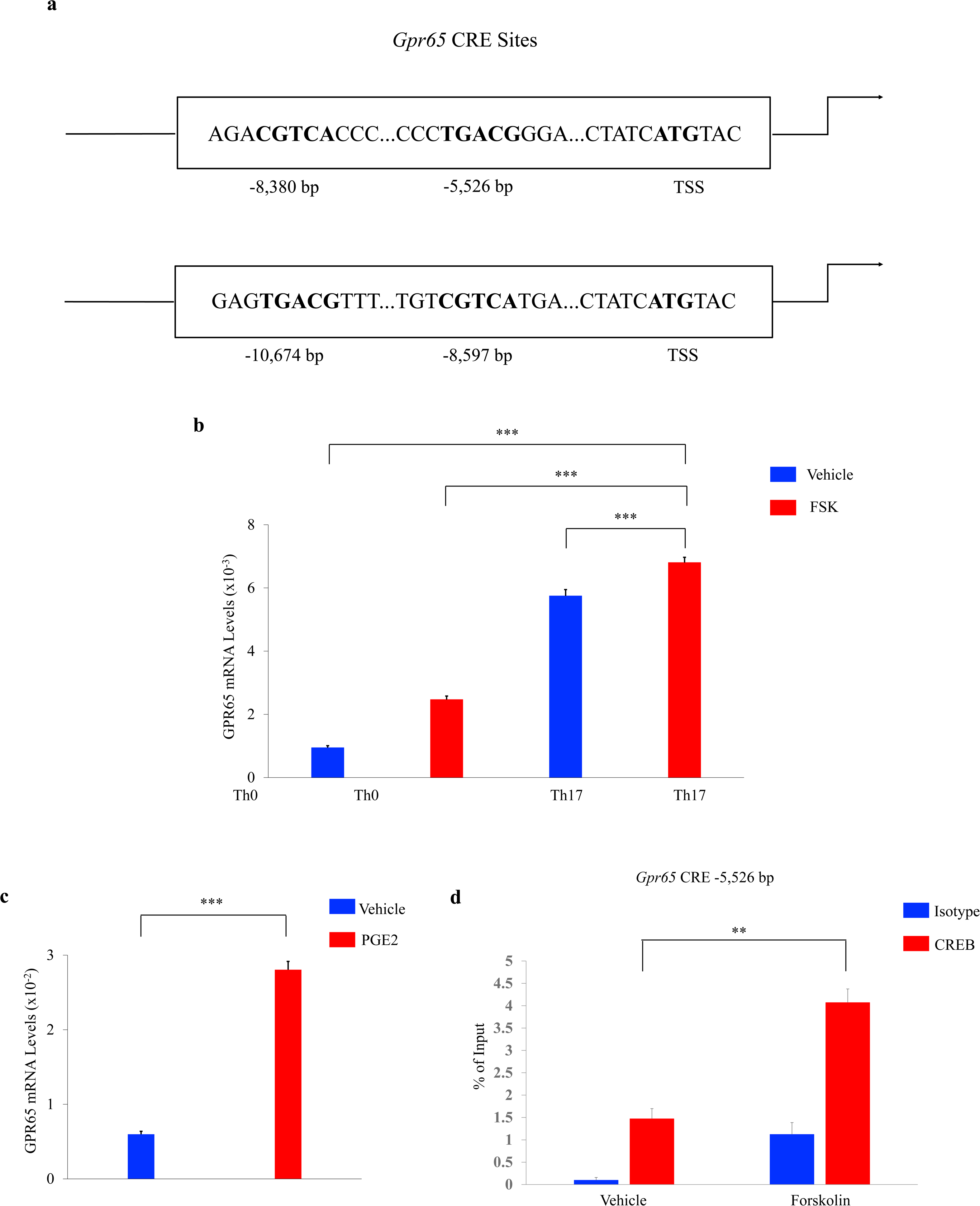
CREB binds upstream of GPR65 transcriptional start site. (a) CRE half-site locations upstream of the mouse GPR65 TSS (NC_000078.7; Chromosome 12; GRCm39; C57BL/6J). (b) GPR65 mRNA levels in naïve CD4⁺ T cells under Th0 or Th17 polarization treated with vehicle (DMSO) or forskolin (10 µM). Vehicle or forskolin was added for 1 hour on day 4 of Th17 differentiation prior to RNA extraction. (c) GPR65 mRNA levels in naïve CD4⁺ T cells under Th17 conditions treated with vehicle (DMSO) or PGE₂ (10 µM) for 1 hour. (d) Chromatin immunoprecipitation (ChIP) showing CREB recruitment upstream of the GPR65 TSS in naïve CD4⁺ T cells cultured under Th17 conditions with vehicle (DMSO) or forskolin (10 µM). Vehicle or forskolin was added for 1 hour on day 4 of Th17 differentiation prior to crosslinking and chromatin preparation. Isotype (IgG) controls were included for both conditions. All bar graphs represent mean ± s.e.m., n = 3 independent experiments. Statistical analyses were performed using unpaired Student’s t-test (*P < 0.05; **P < 0.005; ***P < 0.0005).

To determine if *Gpr65* is transcriptionally regulated by CREB and it is known that CREB is activated in part by increased levels of cAMP, we stimulated differentiated Th17 cells with the cAMP agonist forskolin and measured expression of *Gpr65*. Indeed, exposure to forskolin further increased *Gpr65* expression in Th17 cells (**Fig. 2b**). We have previously shown that the CREB/CRTC2 pathway regulates IL-17A and IL-17F expression by binding to their promoters in response to PGE2 stimulation [13]. Similarly, PGE2 stimulation led to enhanced expression of *Gpr65* in Th17 cells (**Fig. 2c**).

To determine whether *Gpr65* is a direct transcriptional target of CREB, we performed chromatin immunoprecipitation (ChIP) assays. Forskolin stimulation markedly increased CREB binding at the −5,526 CRE half-site (5′-TGACG-3′) within the Gpr65 promoter (Fig. 2d). Isotype (IgG) controls were included for both treatment conditions. Although a modest increase in IgG signal was observed with forskolin, which is consistent with the global chromatin accessibility changes known to occur following cAMP–CREB pathway activation, the increase in CREB enrichment was substantially greater. Together, these data demonstrate that CREB is specifically recruited to the *Gpr65* promoter in response to forskolin stimulation.

### Decreased Th17 cell differentiation in Gpr65^-/-^ mice

To evaluate the role of CREB in regulating *Gpr65*, we expressed a dominant-negative CREB polypeptide called A-CREB under Th17 differentiation conditions. A-CREB expression significantly decreased *Gpr65* expression in Th17 differentiated cells (**Fig. 3a**). Consistent with our previous findings [13] A-CREB expression also decreased IL-17A mRNA levels suggesting a decrease in Th17 cell differentiation (**Fig. 3b**). These in vitro findings reflect effects on Th17 differentiation, not pathogenicity, which is assessed separately in the EAE model

**Figure 3.**
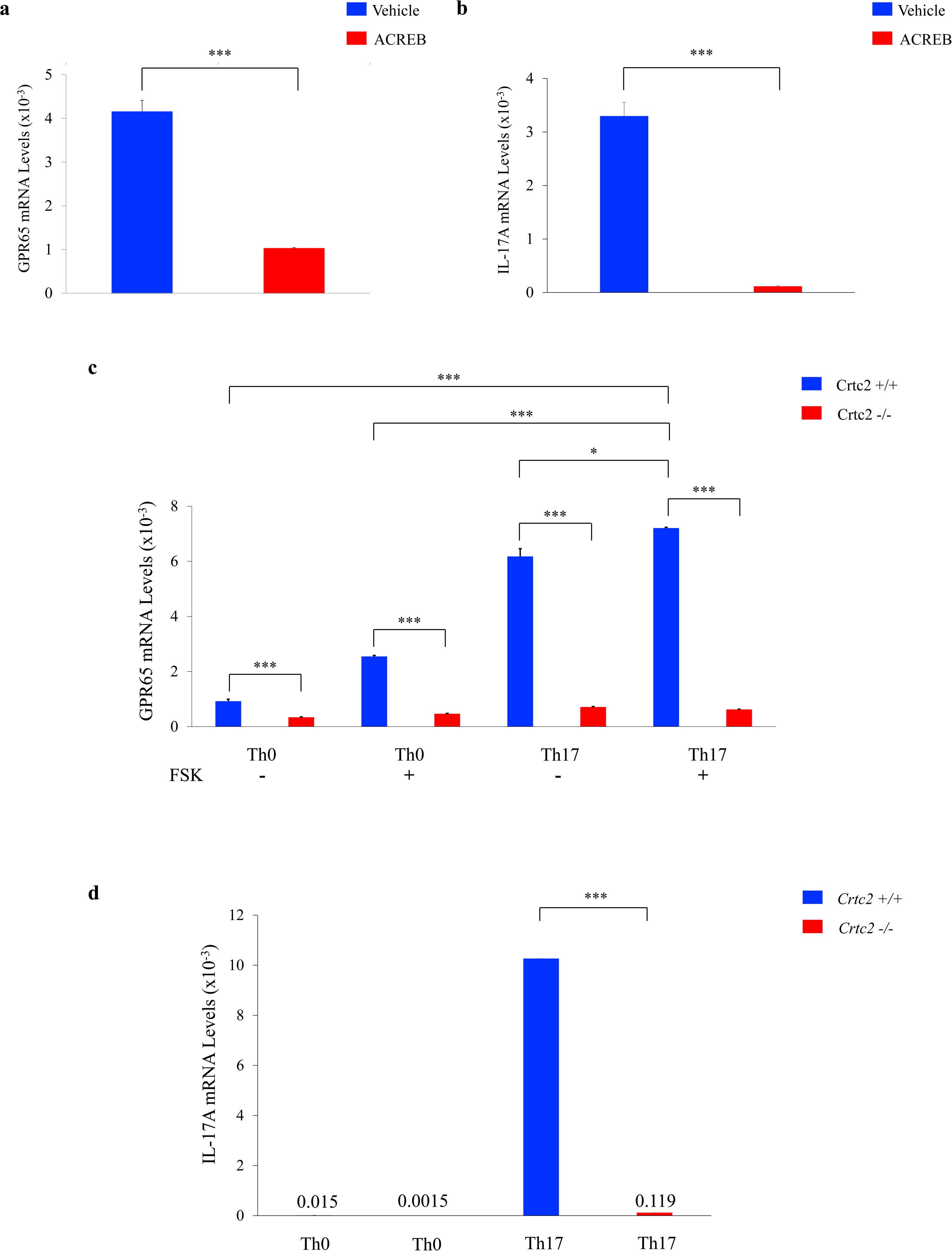
GPR65 regulation by the CRTC2/CREB pathway. (a) Effect of A-CREB expression on GPR65 mRNA levels in naïve CD4⁺ T cells under Th17 differentiation conditions. (b) Effect of A-CREB on IL-17A mRNA levels under Th17 differentiation conditions; Th0 cells included as non-polarized controls. Forskolin (10 µM) added as indicated. Forskolin was added for 1 hour on day 4 of Th17 differentiation prior to RNA extraction. (c) GPR65 mRNA levels in wild-type and CRTC2⁻/⁻ T cells cultured under Th17 conditions. (d) IL-17A mRNA levels in wild-type and CRTC2⁻/⁻ naïve CD4⁺ T cells cultured under Th17 conditions. Data represent mean ± s.e.m., n = 3 independent experiments. Statistical analyses were performed using unpaired Student’s t-test (*P < 0.05; **P < 0.005; ***P < 0.0005).

The CREB co-activator, CRTC2, has been demonstrated to enhance IL-17A and IL-17F mRNA levels in Th17 cells [13]. To determine if CRTC2 enhances *Gpr65* mRNA levels, we differentiated naïve CD4^+^ T cells from *Crtc2^-/-^* and wildtype mice into Th17 cells. As expected, *Gpr65* mRNA levels were decreased in *Crtc2^-/-^*Th17 cells (**Fig. 3c**). We also confirmed a decrease in Th17 cells via IL-17A mRNA levels with loss of CRTC2 (**Fig. 3d**). This loss was not due to decrease in proliferation or survival of Th17 cells since we previously reported no changes in levels of Annexin-V staining or anti-apoptotic factors Bcl-2 and Bcl-xL between wild-type and *Crtc2^-/-^*cells under Th17 differentiation conditions [13]. These results indicate that both CRTC2 and CREB regulate *Gpr65* gene expression by binding upstream of its TSS in response to stimulation.

Previously, the role of GPR65 had only been determined in Th17 cells. To investigate if GPR65 plays a role in other helper T cells subsets, naïve CD4⁺ T cells from whole body *Gpr65^-/-^*mice were differentiated into Th17 cells and regulatory T (Treg) cells, and Th1 cells. There was no statistically significant differences in Th1 cells (**Fig. 4a)** or Treg cell subsets (**Fig. 4b**) between *Gpr65^-/-^* and wildtype differentiated T-cells. However, there was a statistically significant decrease in Th17 cells from *Gpr65^-/-^*differentiated T-cells compared to wildtype differentiated T-cells **(Fig. 4c)**. We also did not detect any differences in IFN-γ or FOXP3 mRNA levels in Th1 and Treg cells, respectively, between *Gpr65^-/-^* and wildtype differentiated cells (Fig. 4d). As expected, we did observe statistically significant decreases in IL-17A levels in *Gpr65^-/-^* differentiated Th17 cells compared to wildtype.

**Figure 4.**
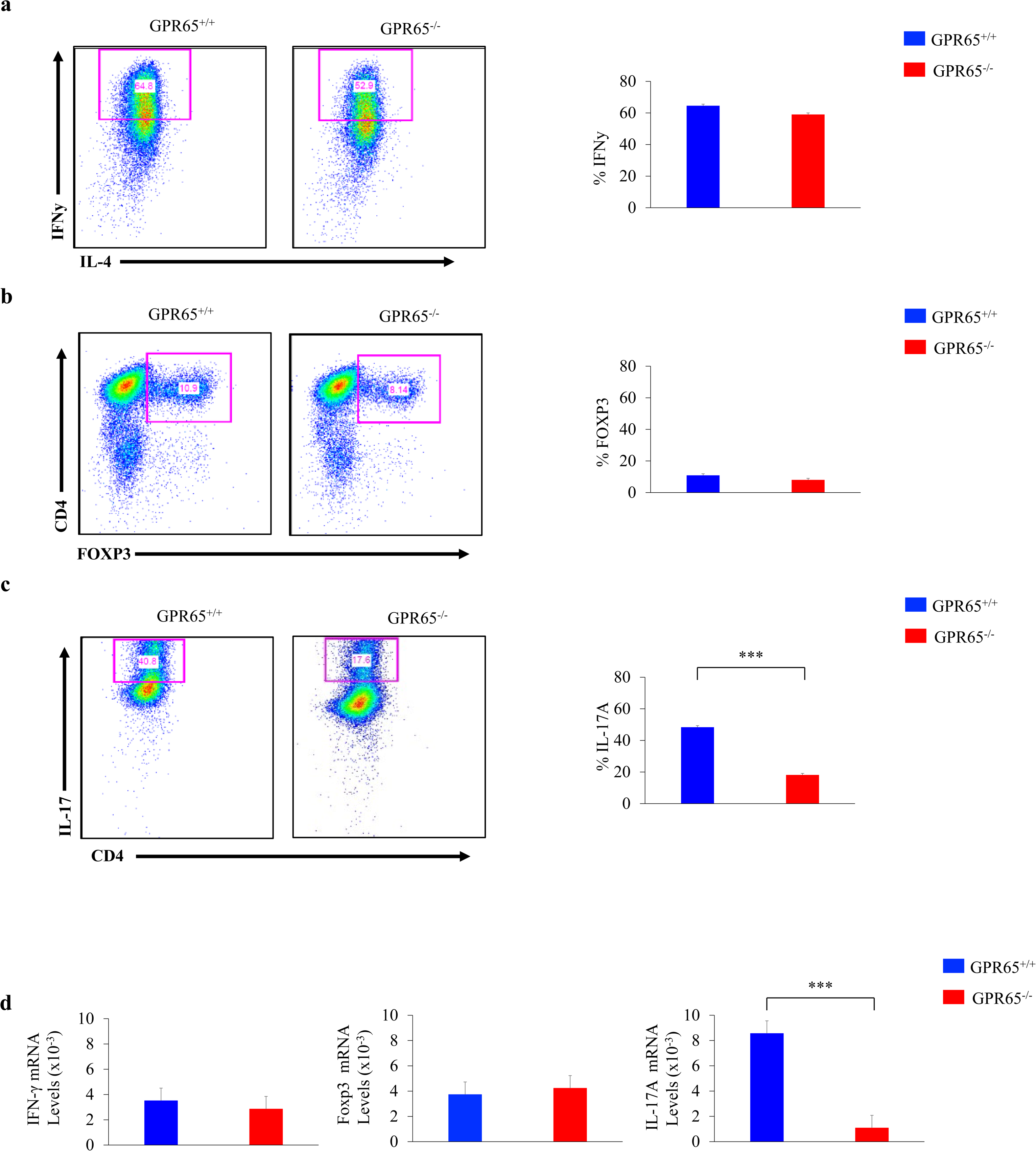
Decreased Th17 cell differentiation in *Gpr65^-/-^* mice. Flow cytometry plots (left) and quantification (right) of naïve CD4⁺ T cells differentiated under (a) Th1, (b) Treg, and (c) Th17 conditions from wild-type and *Gpr65^-/-^* mice. All samples were gated using the following sequence: (1) singlet discrimination (FSC-H × FSC-A), (2) live/dead exclusion, and (3) CD4⁺ T-cell gating, followed by cytokine- or transcription factor–specific gating (IFN-γ, Foxp3, or IL-17A) depending on the differentiation condition. Representative dot plots are shown (left) with corresponding quantification (right). (d) mRNA levels of lineage-specific genes in naïve CD4⁺ T cells differentiated under Th1, Treg, or Th17 conditions from wild-type and *Gpr65^-/-^* mice. Bar graphs represent mean ± s.e.m., n = 3 independent experiments. Statistical analyses were performed using unpaired Student’s t-test (*P < 0.05; **P < 0.005; ***P < 0.0005).

### GPR65 regulates Th17 cell proliferation, survival and metabolism

To determine whether GPR65 influences Th17 cell proliferation and survival, we first examined Annexin V staining relative to CFSE intensity by two-parameter flow cytometry. GPR65^⁻/⁻^ Th17 cells displayed reduced Annexin V⁺ staining compared to wild-type cells, indicating fewer apoptotic events at the time of analysis (**Fig. 5a**). Because Th17 cultures typically undergo extensive and asynchronous divisions, CFSE profiles often do not resolve into discrete division peaks [2].

**Figure 5.**
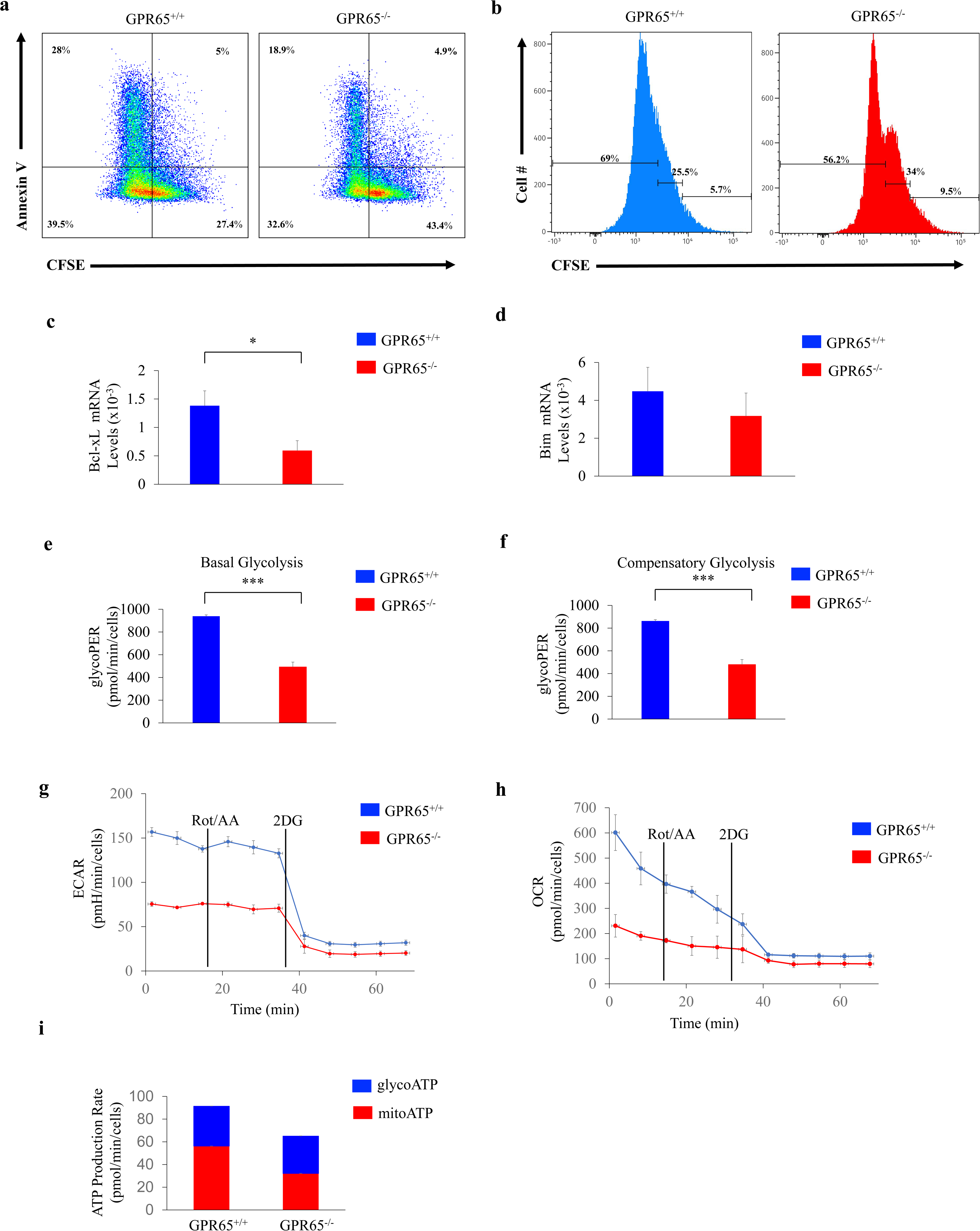
GPR65 regulates Th17 cell survival, proliferation and metabolism. (a) Representative flow cytometry plots of Annexin V versus CFSE staining in wild-type and *Gpr65^⁻/⁻^* naïve CD4⁺ T cells differentiated under Th17 conditions. *Gpr65^⁻/⁻^* Th17 cells show fewer Annexin V⁺ cells within the proliferating population relative to wild type. (b) CFSE histograms of wild-type and *Gpr65^⁻/⁻^* Th17-differentiated cells, demonstrating reduced CFSE dilution in *Gpr65^⁻/⁻^* cultures. CFSE intensity was divided into CFSE^hi^ (undivided), CFSE^int^ (partially divided), and CFSE^lo^ (extensively divided) gates to quantify proliferative progression. (c) Bcl-xL mRNA levels and (d) Bim mRNA levels in wild-type and *Gpr65^-/-^*Th17 cells. (e) basal glycolysis, (f) compensatory glycolysis, (g) extracellular acidification rate (ECAR), and (h) oxygen consumption rate (OCR) and (i) ATP production rate measured by Seahorse analysis in wild-type and *Gpr65^-/-^*Th17 cells. Data represent mean ± s.e.m., n = 3 independent experiments. Statistical analyses were performed with unpaired Student’s t-test (*P < 0.05; **P < 0.005; ***P < 0.0005).

Consistent with this, the wild-type Th17 population showed a broad CFSE-low profile, whereas an intermediate CFSE population became readily distinguishable only in the GPR65^⁻/⁻^ cultures. While the Annexin V versus CFSE plot suggested differences in CFSE intensity between genotypes, the CFSE histogram (Fig. 5b) revealed three distinct CFSE intensity ranges, prompting us to quantify proliferative progression using CFSE^hi^ (undivided), CFSE intermediate (CFSE^int^; partially divided), and CFSE^lo^ (extensively divided) gates. Compared with wild-type Th17 cells, GPR65^⁻/⁻^ cells accumulated within the CFSE^int^ population and showed a reduced transition into the CFSE^lo^ gate, demonstrating impaired proliferative progression. Additional quantification of CFSE^hi^, CFSE^int^, and CFSE^lo^ subsets, as well as Annexin V staining within each population, is provided in Supplemental Figure 2 (**Fig. S2**). Because fewer *Gpr65^-/-^* cells reached the highly divided CFSE^lo^ state, where apoptosis is typically increased, the overall decrease in Annexin V⁺ cells in *Gpr65^-/-^* cultures reflects reduced proliferative expansion rather than enhanced survival. Thus, the dominant effect of GPR65 deficiency is a defect in Th17 proliferative progression, accompanied by proportionally fewer Annexin V⁺ cells due to limited entry into late-division states.

To further assess survival pathways, we measured expression of apoptosis-associated genes. Bcl-xL mRNA levels were significantly reduced in *Gpr65^-/-^*Th17 cells, whereas Bim mRNA levels showed a modest but non-significant decrease (**Fig. 5c and 5d**). Collectively, these data indicate that GPR65 supports Th17 cell expansion, and that its loss results in reduced progression through proliferative divisions accompanied by changes in Bcl-xL expression.

To investigate the role of GPR65 in metabolism, seahorse metabolic assays were performed on wildtype and *Gpr65^-/-^* differentiated Th17 cells. *Gpr65^-/-^* Th17 cells exhibited decreased basal glycolysis compared to wildtype Th17 cells (**Fig. 5e**). When Th17 cells were treated with rotenone and antimycin, which inhibit mitochondrial respiration and force reliance on glycolysis, *Gpr65^-/-^* Th17 cells were less able to compensate relative to wild-type Th17 cells (**Fig. 5f**). Consistent with impaired glycolysis, extracellular acidification rate (ECAR) was reduced in *Gpr65^-/-^* Th17 cells (**Fig. 5g**). ECAR reflects proton production during glycolysis and serves as a real-time readout of glycolytic flux.

We next assessed whether GPR65 deficiency affects mitochondrial function. *Gpr65^-/-^* Th17 cells displayed a reduction in oxygen consumption rate (OCR), indicating impaired mitochondrial respiration (**Fig. 5h**). Consistent with their reduced glycolytic capacity, GPR65^⁻/⁻^ Th17 cells also showed decreased ATP production (**Fig. 5i**). These findings indicate that GPR65 is required to support the metabolic program necessary for sustained Th17 proliferative progression. Rather than directly regulating survival, GPR65 promotes the bioenergetic capacity that enables Th17 cells to complete multiple rounds of division.

### GPR65 small molecules modulate Th17 cells

Due to GPR65 being highly expressed in Th17 cells, and because loss of GPR65 results in reduced EAE sensitivity [2], targeting GPR65 with small molecules may represent a promising strategy for treating Th17-mediated diseases such as multiple sclerosis. Onozawa et al. identified the GPR65-specific agonist BTB09089 by screening an in-house compound library using a cAMP-based functional assay [18]. More recently, Huang et al. described a negative allosteric inhibitor of BTB09089, Zinc62678696, identified by treating human embryonic kidney 293T cells transfected with GPR65 and observing a reduction in cAMP production [22].

We evaluated the effects of these GPR65-specific small molecules under Th17-differentiation conditions. The GPR65 agonist BTB09089 induced a significant increase in IL-17A mRNA expression (**Fig. 6a**), whereas the GPR65 antagonist Zinc62678696 caused a marked decrease in IL-17A expression (**Fig. 6b**). These findings demonstrate that pharmacologic modulation of Th17 cells is feasible through small-molecule targeting of GPR65.

**Figure 6.**
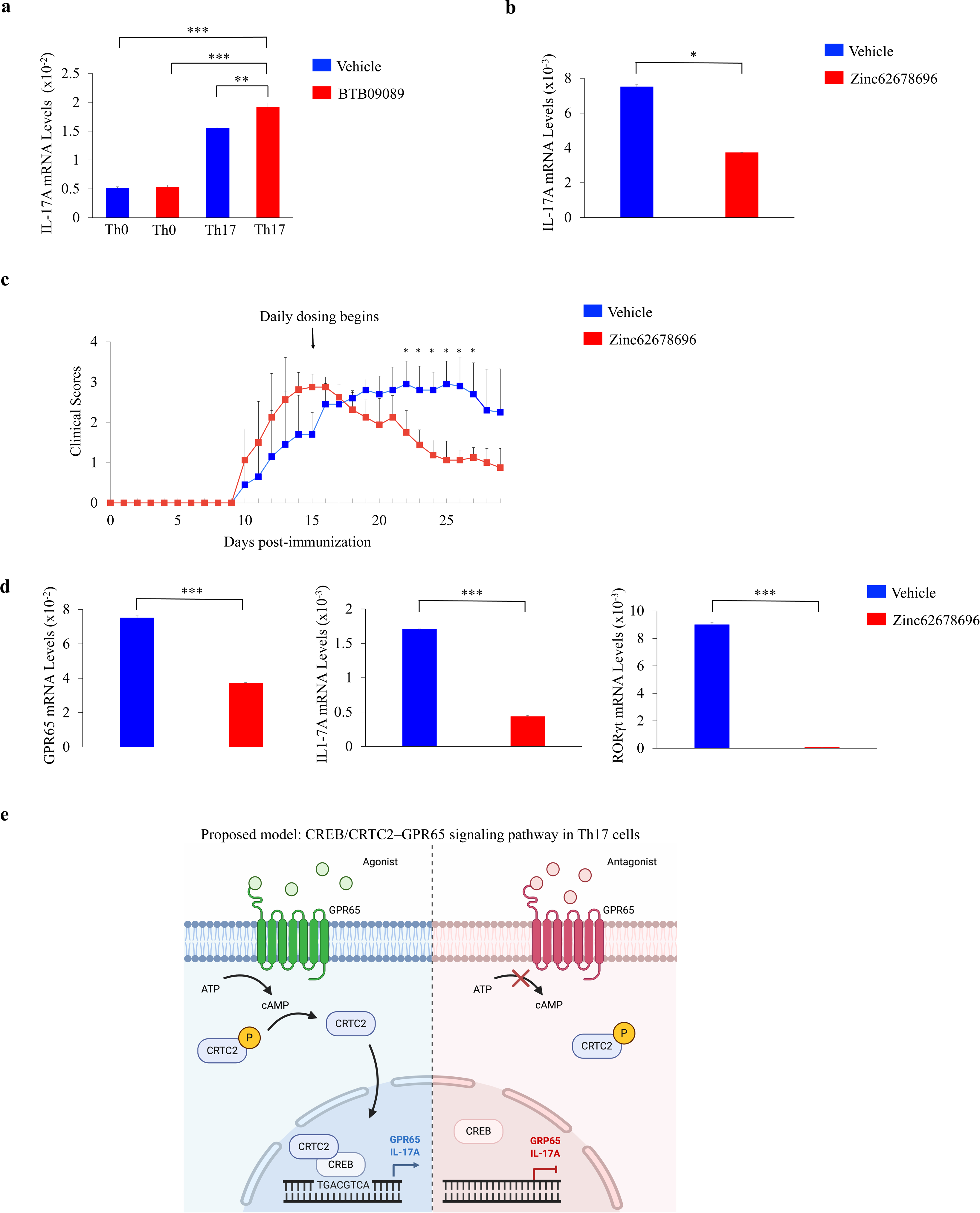
GPR65 small molecules modulate Th17 cells. (a) IL-17A mRNA levels in naïve CD4⁺ T cells differentiated under Th17 conditions with vehicle or BTB09089 (30 µM). Th0 cells are shown as control. BTB09089 was added for 1 hour on day 4 of Th17 differentiation prior to RNA extraction. (b) IL-17A mRNA levels in naïve CD4⁺ T cells under Th17 conditions treated with vehicle (DMSO) or Zinc62678696 (30 µM). Vehicle or Zinc62678696 was added for 1 hour on day 4 of Th17 differentiation prior to RNA extraction. (c) EAE clinical scores of wild-type (n = 10) and *Gpr65^-/-^* (n = 10) mice immunized with MOG₃₅–₅₅. Arrow indicates daily administration of vehicle or Zinc62678696 (75 mg/kg, intraperitoneally, every 24 hours) beginning on day 15 post-immunization. Each line represents the mean clinical score. Daily clinical scores were compared between groups using the non-parametric Mann–Whitney U test, which is appropriate for ordinal EAE scoring. *p* < 0.05 was considered significant. (d) GPR65, IL-17A, and RORγt mRNA levels in human Th17 cells treated with vehicle (DMSO) or Zinc62678696 (30 µM). Vehicle or Zinc62678696 was added for 1 hour prior to RNA extraction. **(e)** Schematic model summarizing CREB/CRTC2-dependent regulation of GPR65 in Th17 cells and the effects of pharmacologic inhibition on Th17 effector function and EAE severity. Activation of GPR65 increases intracellular cAMP and promotes phosphorylation of CREB and dephosphorylation/nuclear translocation of CRTC2. The CREB/CRTC2 complex enhances transcription of *Il17a* and also induces *Gpr65*, forming a positive feedback loop that reinforces GPR65 signaling and sustains Th17 effector function. Pharmacologic inhibition of GPR65 disrupts this signaling axis, resulting in reduced IL-17A expression in vitro and decreased Th17-mediated pathology in vivo, including attenuation of EAE severity. Data in (a), (b), and (d) represent mean ± s.e.m. from n = 3 independent experiments. Statistical analyses were performed using unpaired Student’s t-test (*P < 0.05; **P < 0.005; *P < 0.0005), unless otherwise indicated.

To assess whether pharmacologic blockade of GPR65 affects Th17 pathogenicity in vivo, we induced EAE in wild-type mice and initiated Zinc62678696 treatment at symptom onset. Beginning on day 15 post-immunization, mice received daily administration of Zinc62678696, and clinical scores were monitored thereafter. Zinc62678696-treated mice exhibited a clear reduction in EAE severity compared with vehicle-treated controls (**Fig. 6c**). These findings indicate that pharmacologic inhibition of GPR65 is sufficient to attenuate Th17-mediated pathology in vivo.

Lastly, to determine whether GPR65 also contributes to human Th17 cell function, we treated human Th17 cells with Zinc62678696. As anticipated, treatment resulted in a significant decrease in GPR65, IL-17A, and RORɣt mRNA levels compared with vehicle-treated cells (**Fig. 6d**). These results collectively indicate that pharmacologic blockade of GPR65 using small-molecule antagonists can suppress both mouse and human Th17 cell responses.

## DISCUSSION

Here, we demonstrate for the first time that the orphan G protein–coupled receptor GPR65 is transcriptionally regulated by the CRTC2/CREB pathway. We previously showed that loss of GPR65 in T cells reduces Th17 differentiation and protects mice in the experimental autoimmune encephalomyelitis (EAE) model of multiple sclerosis [13]. GPR65 has additionally been linked to multiple sclerosis, ankylosing spondylitis, inflammatory bowel disease, and Crohn’s disease [23–25], suggesting that understanding its regulation may be important for targeting Th17-mediated inflammatory disorders. Our in vitro data specifically addresses Th17 differentiation, whereas the EAE experiments demonstrate effects on Th17 pathogenicity, a distinct functional property of Th17 cells that drives autoimmune tissue damage.

Our RNA-seq analysis identified *Gpr65* as a CREB-responsive gene in Th17 cells treated with forskolin, implicating cAMP signaling in its transcriptional control. We found that GPR65 is highly expressed in Th17 cells compared with other CD4⁺ T-cell subsets, further supporting its functional importance in this lineage. Two additional GPCR genes, *Gpr18* and *Gpr132*, were also enriched in Th17 cells in our dataset, indicating that other GPCRs may contribute to Th17 biology and warrant future investigation. Notably, expression of *Gpr65* was markedly reduced in CD8⁺ T cells, suggesting that GPR65 may play a more specialized role in CD4⁺ T-cell function.

Chromatin immunoprecipitation revealed CREB binding to an upstream enhancer region of *Gpr65* located 5,526 bp upstream of the transcriptional start site (TSS), consistent with prior studies showing that CRE sites can reside in either promoter or enhancer elements [12]. Forskolin treatment increased *Gpr65* expression in wild-type Th17 cells but not in CRTC2^⁻/⁻^ Th17 cells, indicating that CRTC2 is required for CREB-mediated induction of GPR65. Similarly, overexpression of the dominant-negative CREB inhibitor, A-CREB, suppressed *Gpr65* expression. Together, these findings support a model in which cAMP elevation activates the CREB/CRTC2 pathway to upregulate GPR65 expression in Th17 cells.

Gaublomme et al. previously reported that GPR65 deficiency reduces Th17 differentiation by approximately 40% [2], consistent with our observations that whole-body GPR65 knockout decreases Th17 output. To understand the mechanism underlying this reduction, we examined proliferation, apoptosis, and metabolism in GPR65-deficient Th17 cells. Two-parameter flow cytometry revealed that *Gpr65^-/-^* Th17 cells exhibited reduced Annexin V⁺ staining, but this decrease reflected the fewer cells reaching late CFSE^lo^ divisions, where apoptosis normally increases, rather than enhanced survival. CFSE profiling showed that *Gpr65^-/-^* Th17 cells accumulated in the CFSE^int^ population and failed to efficiently progress into the extensively divided CFSE^lo^ gate, demonstrating a defect in sustained proliferative progression. Although *Gpr65^-/-^* Th17 cells expressed lower levels of the pro-survival factor Bcl-xL, this reduction likely reflects impaired metabolic fitness rather than a primary defect in apoptotic regulation.

Our metabolic analyses support this interpretation. *Gpr65^-/-^*Th17 cells displayed reduced basal glycolysis, decreased extracellular acidification rate (ECAR), diminished compensatory glycolysis under mitochondrial inhibition, and lower oxygen consumption rate (OCR). Together, these defects result in significantly reduced ATP production from both glycolysis and oxidative phosphorylation. Because Th17 cells rely heavily on glycolytic and mitochondrial metabolism to fuel multiple rounds of division, this impaired metabolic program provides a mechanistic explanation for the failure of *Gpr65^-/-^* cells to complete proliferative progression. Thus, rather than directly regulating differentiation or survival, GPR65 promotes the metabolic fitness required to sustain Th17 proliferation, and its loss restricts Th17 expansion by limiting bioenergetic capacity.

Current therapies for Th17-mediated diseases, such as multiple sclerosis, rely on broad immunosuppression, which can increase susceptibility to opportunistic infections. In this study, we identify the small-molecule antagonist Zinc62678696 as a selective inhibitor of GPR65 signaling. In vitro, the GPR65 agonist BTB09089 enhanced IL-17A expression in Th17 cells, whereas Zinc62678696 markedly reduced IL-17A expression. In the EAE model, therapeutic administration of Zinc62678696 beginning at symptom onset led to a noticeable reduction in disease severity relative to vehicle controls. The therapeutic benefit observed when treatment was initiated at symptom onset likely reflects the ability of GPR65 blockade to rapidly suppress IL-17A transcription and diminish the metabolic fitness of effector Th17 cells already infiltrating the CNS. Importantly, these effects translated to human cells, as Zinc62678696 treatment decreased expression of GPR65, IL-17A, and RORγt in human Th17 cells. The rapid transcriptional decreases observed after brief GPR65 inhibition likely reflect early disruption of signaling inputs that sustain ongoing *RORC* and *IL17A* transcription, rather than immediate remodeling of the Th17 transcriptional program.

These data support a model in which CREB/CRTC2-dependent induction of *Gpr65* enhances Th17 metabolic fitness and effector function, positioning GPR65 as a central regulator of Th17 pathogenicity (**Fig. 6e**). GPR65 signals through a canonical G_s_–cAMP pathway that has been extensively characterized in multiple cell types. Elevated cAMP activates PKA, which phosphorylates CREB and promotes the dephosphorylation and nuclear translocation of CRTC2—events that are well established across numerous biochemical studies. Because these signaling steps are not unique to Th17 cells, our work focuses on how CREB/CRTC2-dependent transcriptional induction of Gpr65 integrates into the Th17 effector program. This pathway provides a mechanistic basis for how CREB/CRTC2 activation and GPR65 signaling reinforce one another to sustain *Il17* transcription and Th17 effector function in inflammatory environments.

Together, our findings reveal that GPR65 is a cAMP-responsive, CREB/CRTC2-regulated effector that promotes Th17 cell metabolism and survival. In this framework, pharmacologic inhibition of GPR65 disrupts this signaling axis, resulting in reduced IL-17A expression in vitro and attenuated disease severity in vivo. Targeting GPR65 may therefore represent a promising strategy for selectively suppressing pathogenic Th17 responses while preserving broader immune function. Future studies may also explore monoclonal approaches to GPR65 blockade as complementary modalities to expand therapeutic options for Th17-mediated autoimmune disease.

## ACKNOWLEDGEMENTS

We thank Dr. Marc Montminy of the Salk Institute for his thoughtful input, lab space and resources to conduct this research. At UC Irvine, we would like to thank Dr. Craig Walsh for his invaluable guidance and allowing us to use his animal lab space, Julio Alejandro Ayala Angulo for his technical advice and the Stem Cell Research Center Flow and Mass Cytometry Core facility, specifically Vanessa Scarfone and Pauline Nguyen. This work was supported by the Joseph H. Stahlberg Foundation.

