## Supplementary Figure 1 for "CREB/CRTC2-Induced GPR65 Orchestrates Th17 Metabolic Fitness and Pathogenic Effector Function"

### Slide 1
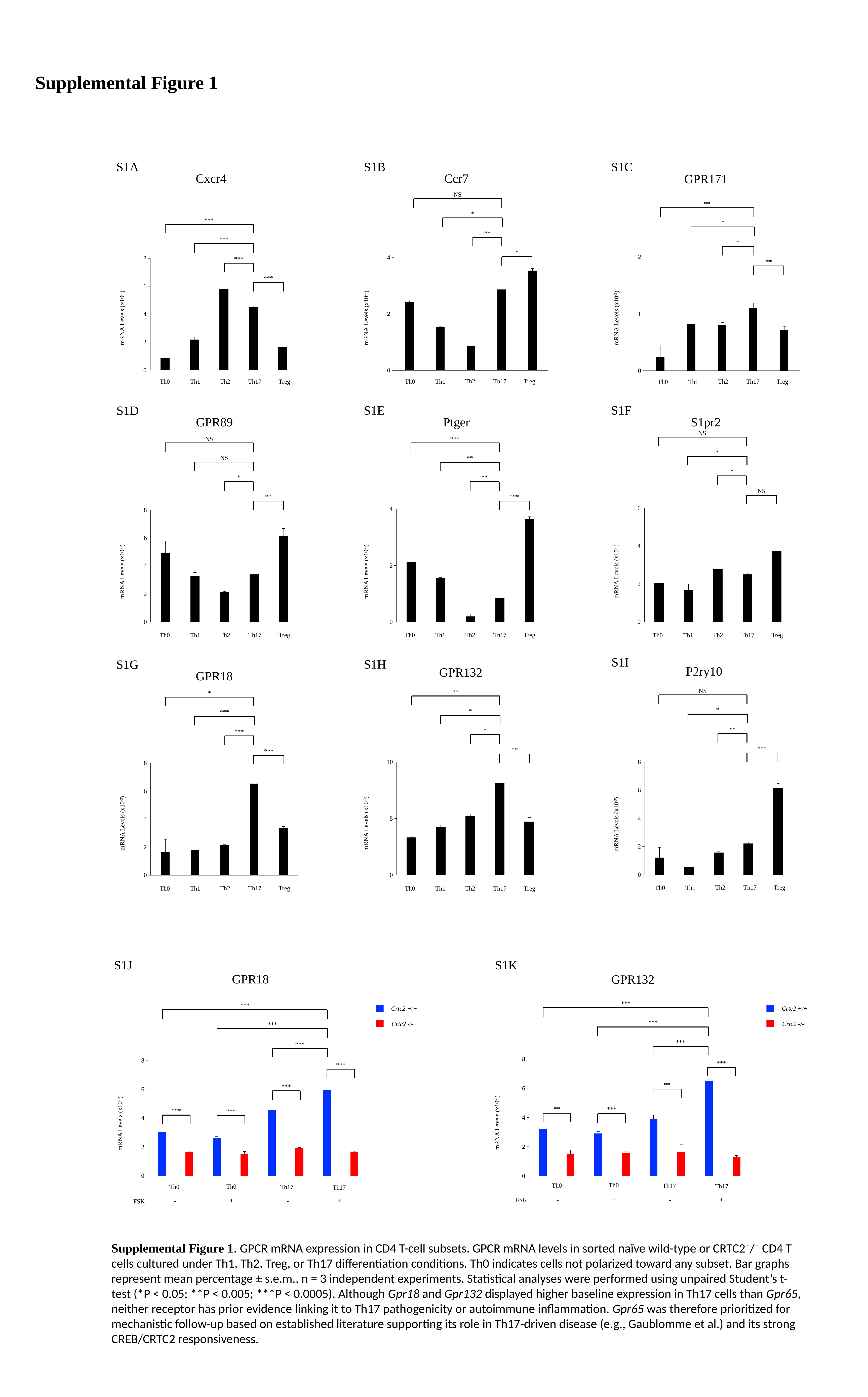

Supplemental Figure 1
S1A
S1B
S1C
Ccr7
Cxcr4
GPR171
NS
**
*
***
*
**
***
*
#### Chart
| Category | |
|---|---|
| Th0 | 0.238139755035311 |
| Th1 | 0.82294325163455 |
| Th2 | 0.80035137753343 |
| Th17 | 1.09989155803602 |
| Treg | 0.708377164649693 |
#### Chart
| Category | |
|---|---|
| Th0 | 2.40724222190264 |
| Th1 | 1.5339903796121 |
| Th2 | 0.873165171524453 |
| Th17 | 2.86271693504259 |
| Treg | 3.53261865626585 |
#### Chart
| Category | |
|---|---|
| Th0 | 0.838807910623503 |
| Th1 | 2.18218808662711 |
| Th2 | 5.8132047773814 |
| Th17 | 4.47703429453255 |
| Treg | 1.65466044434139 |*
***
**
***
mRNA Levels (x10^-3^)
mRNA Levels (x10^-2^)
mRNA Levels (x10^-2^)
Treg
Th2
Th17
Treg
Th0
Th1
Th2
Th17
Treg
Th0
Th1
Th2
Th17
Th0
Th1
S1F
S1E
S1D
Ptger
S1pr2
GPR89
NS
NS
***
*
NS
**
*
*
**
NS
**
***
#### Chart
| Category | |
|---|---|
| Th0 | 2.02754422541981 |
| Th1 | 1.65220806251129 |
| Th2 | 2.80634318964626 |
| Th17 | 2.49912647786485 |
| Treg | 3.74948919805386 |
#### Chart
| Category | |
|---|---|
| Th0 | 2.12058437212102 |
| Th1 | 1.55918589975053 |
| Th2 | 0.187715766789612 |
| Th17 | 0.847825651704341 |
| Treg | 3.64526099291683 |
#### Chart
| Category | |
|---|---|
| Th0 | 4.93127 |
| Th1 | 3.25617 |
| Th2 | 2.11909 |
| Th17 | 3.39456 |
| Treg | 6.14691 |mRNA Levels (x10^-3^)
mRNA Levels (x10^-4^)
mRNA Levels (x10^-5^)
Treg
Th2
Th17
Treg
Th0
Th1
Treg
Th2
Th17
Th2
Th17
Th0
Th1
Th0
Th1
S1I
S1H
S1G
P2ry10
GPR132
GPR18
NS
**
*
*
*
***
**
*
***
***
**
#### Chart
| Category | |
|---|---|
| Th0 | 1.19638124249805 |
| Th1 | 0.535940693684654 |
| Th2 | 1.55149585099837 |
| Th17 | 2.19895378441688 |
| Treg | 6.10789521940829 |
#### Chart
| Category | |
|---|---|
| Th0 | 3.308938 |
| Th1 | 4.200082 |
| Th2 | 5.169326 |
| Th17 | 8.123374 |
| Treg | 4.720051 |***
#### Chart
| Category | |
|---|---|
| Th0 | 1.62912300147238 |
| Th1 | 1.791034278178 |
| Th2 | 2.14740917347217 |
| Th17 | 6.53170491677407 |
| Treg | 3.38556388889341 |mRNA Levels (x10^-3^)
mRNA Levels (x10^-3^)
mRNA Levels (x10^-3^)
Treg
Th2
Th17
Th0
Th1
Treg
Th2
Th17
Th0
Th1
Treg
Th2
Th17
Th0
Th1
S1J
S1K
GPR18
GPR132
***
***
Crtc2 +/+
Crtc2 +/+
***
Crtc2 -/-
Crtc2 -/-
***
#### Chart
| Category | |
|---|---|***
***
#### Chart
| Category | |
|---|---|***
***
**
***
**
***
***
***
mRNA Levels (x10^-3^)
mRNA Levels (x10^-3^)
 Th0
Th0
Th17
Th17
 Th0
Th0
Th17
Th17
 - + - +
FSK
 - + - +
FSK
Supplemental Figure 1. GPCR mRNA expression in CD4 T-cell subsets. GPCR mRNA levels in sorted naïve wild-type or CRTC2⁻/⁻ CD4 T cells cultured under Th1, Th2, Treg, or Th17 differentiation conditions. Th0 indicates cells not polarized toward any subset. Bar graphs represent mean percentage ± s.e.m., n = 3 independent experiments. Statistical analyses were performed using unpaired Student’s t-test (*P < 0.05; **P < 0.005; ***P < 0.0005). Although Gpr18 and Gpr132 displayed higher baseline expression in Th17 cells than Gpr65, neither receptor has prior evidence linking it to Th17 pathogenicity or autoimmune inflammation. Gpr65 was therefore prioritized for mechanistic follow-up based on established literature supporting its role in Th17-driven disease (e.g., Gaublomme et al.) and its strong CREB/CRTC2 responsiveness.
