## Supplementary Figure 2 for "CREB/CRTC2-Induced GPR65 Orchestrates Th17 Metabolic Fitness and Pathogenic Effector Function"

### Slide 1
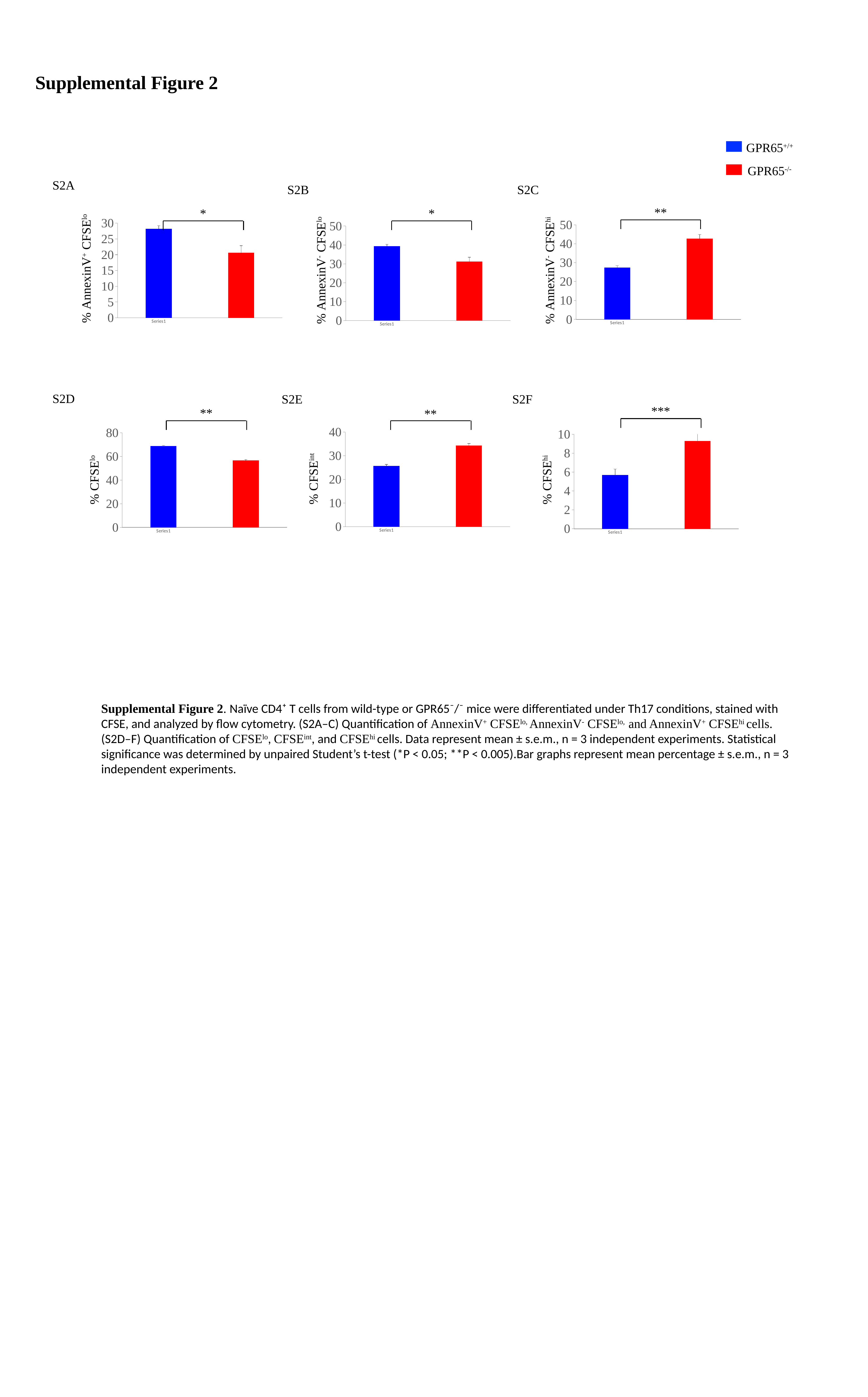

Supplemental Figure 2
GPR65^+/+^
GPR65^-/-^
S2A
S2B
S2C
**
*
*
#### Chart
| Category | Annexin V+ CFSElo |
|---|---|
| | 28.2 |
| | 20.599999999999998 |
#### Chart
| Category | Annexin V- CFSEhi |
|---|---|
| | 27.366666666666664 |
| | 42.56666666666667 |
#### Chart
| Category | Annexin V- CFSElo |
|---|---|
| | 39.3 |
| | 31.2 |% AnnexinV^+^ CFSE^lo^
% AnnexinV^-^ CFSE^lo^
% AnnexinV^-^ CFSE^hi^
S2D
S2E
S2F
***
**
**
#### Chart
| Category | CFSE INT |
|---|---|
| | 25.666666666666668 |
| | 34.233333333333334 |
#### Chart
| Category | CFSElo |
|---|---|
| | 68.66666666666667 |
| | 56.46666666666667 |
#### Chart
| Category | CFSEhi |
|---|---|
| | 5.666666666666667 |
| | 9.266666666666667 |% CFSE^lo^
% CFSE^int^
% CFSE^hi^
Supplemental Figure 2. Naïve CD4⁺ T cells from wild-type or GPR65⁻/⁻ mice were differentiated under Th17 conditions, stained with CFSE, and analyzed by flow cytometry. (S2A–C) Quantification of AnnexinV^+^ CFSE^lo,^ AnnexinV^-^ CFSE^lo,^ and AnnexinV^+^ CFSE^hi^ cells.
(S2D–F) Quantification of CFSE^lo^, CFSE^int^, and CFSE^hi^ cells. Data represent mean ± s.e.m., n = 3 independent experiments. Statistical significance was determined by unpaired Student’s t-test (*P < 0.05; **P < 0.005).Bar graphs represent mean percentage ± s.e.m., n = 3 independent experiments.
